# The coral symbiont *Candidatus* Aquarickettsia is variably abundant in threatened Caribbean acroporids and transmitted horizontally

**DOI:** 10.1101/2021.01.28.428674

**Authors:** Lydia J. Baker, Hannah G. Reich, Sheila A. Kitchen, J. Grace Klinges, Hanna R. Koch, Iliana B. Baums, Erinn Muller, Rebecca Vega Thurber

## Abstract

The aquatic symbiont “*Candidatus Aquarickettsia rohweri*” infects a diversity of non-bilaterian metazoan phyla. In the threatened coral *Acropora cervicornis*, *Aquarickettsia* proliferates in response to increased nutrient exposure, resulting in suppressed growth and increased disease susceptibility and mortality. This study evaluated the extent, as well as the ecology and evolution of *Aquarickettsia* infecting the Caribbean corals: *Ac. cervicornis* and *Ac. palmata* and their hybrid (‘*Ac. prolifera*’). The bacterial parasite *Aquarickettsia* was found in all acroporids, with host and sampling location impacting infection magnitude. Phylogenomic and genome-wide single nucleotide variant analysis found *Aquarickettsia* clustering by region, not by coral taxon. Fixation analysis suggested within coral colonies, *Aquarickettsia* are genetically isolated to the extent that reinfection is unlikely. Relative to other Rickettsiales, *Aquarickettsia* is undergoing positive selection, with Florida populations experiencing greater positive selection relative to the other Caribbean locations. This may be due to *Aquarickettsia* response to increased nutrient stress in Florida, as indicated by greater *in situ* replication rates in these corals. *Aquarickettsia* did not significantly codiversify with either coral animal nor algal symbiont, and qPCR analysis of gametes and juveniles from susceptible coral genotypes indicated absence in early life stages. Thus, despite being an obligate parasite, *Aquarickettsia* must be horizontally transmitted via coral mucocytes, an unidentified secondary host, or a yet unexplored environmentally mediated mechanism. Importantly, the prevalence of *Aquarickettsia* in *Ac. cervicornis* and high abundance in Florida populations suggests that disease mitigation efforts in the US and Caribbean should focus on preventing early infection via horizontal transmission.

## Introduction

The alpha-proteobacterium “*Ca*. Aquarickettsia *rohweri*” is a symbiont that infects a diversity of aquatic non-bilaterian metazoan phyla from around the world [1]. This includes reef-building corals (scleractinians) (51), as well as other cnidarians (11), sponges (76), kelp mucus, and ctenophores [1]. Although fairly ubiquitous, *A. rohweri* may have a more pervasive interaction with Caribbean acroporids, as Rickettsiales-like organisms, likely to be *A. rohweri,* have been found in all histological examinations of these coral species since 1975 [2–5]. The genome of *A. rohweri* associated with Caribbean acroporids is significantly reduced (1.28 Mbp) and has limited metabolic capacities, including the inability to produce multiple amino acids and ATP [1]. Thus, *A. rohweri* is likely an obligate symbiont dependent on its host for nutrition and energy rather than free-living, but its transmission routes are not yet known. Within Caribbean acroporids, *A. rohweri* have been found in high concentrations in *Acropora cervicornis* exposed to rising nutrient concentrations [6] and in disease-susceptible genotypes [7] and *A. rohweri* has a possible role in the progression of white band disease (WBD) [6, 8, 9]. WBD has contributed significantly to the decline of the reef-building corals, *Ac. cervicornis,* and to a lesser extent, *Ac. palmata* [10, 11]. These two corals are now so rare that they have been listed as threatened under the US Endangered Species Act [12, 13]. Worldwide, coral diseases have contributed to regional losses of between 5-80% of coral cover [10, 14]. Therefore, it is of broad interest to gain a better understanding of how this parasitic symbiont is evolving and transmitted to inform disease management.

Transmission mode is a major determinant of symbiont population structure and evolution [15–17]. Symbionts can be transmitted either vertically via direct transfer from one host generation to the next, or horizontally from a secondary host or the environment. Vertically transmitted symbiont phylogenies are congruent with their host phylogenies, as has been observed in insects [17, 18] and inferred in deep sea clam symbionts [19, 20] and sponge symbionts [21]. Many Rickettsiales species closely related to *A. rohweri* are known to be transmitted vertically, including Wolbachia [16, 22]. In the absence of significant codiversification, symbionts are likely transmitted through an alternative host or through the environment [23, 24]. Close-proximity environmental transmission has been posited for symbionts associated with insect and nematode species [15] as well as flashlight fish [25, 26]. Because *A. rohweri* associates with evolutionarily distant hosts [1], this parasite may be transmitted horizontally similar to arthropod or plant mediated transmission of terrestrial Rickettsiales [27, 28]. Although secondary hosts have not been identified yet, possible modes of transmission include the gastropod *Coralliophila abbreviata* [29], zooplankton [30], or other coral associates. In fact, microscopy of infected coral polyps cannot resolve whether *A. rohweri* is associated with the coral animal or the coral’s obligate, intra-cellular algal mutualist, Symbiodiniaceae. Rickettsiales-like organisms were observed in the actinopharynx, cnidoglandular bands, gastrodermal mucocytes, oral disk, and tentacles of a healthy *Ac. cervicornis* [8], spaces also shared by the algal mutualist. All Caribbean acroporid species take up their algal symbionts anew from the environment upon larval settlement [31], and thus algal symbionts, and perhaps *A. rohweri* with them, are horizontally transmitted.

Regardless of transmission mode, *A. rohweri* populations may also be structured by host species and the environment, although the latter is difficult to disentangle, as one likely co-varies with the other [32]. For example, the location where the first genome of *A. rohweri* was characterized, the Florida Keys, has been exposed to increasing anthropogenic inputs [33, 34] and wide-spread bleaching events [35, 36]. Coral in this area have also experienced multi-year epizootics, including stony coral tissue loss, white-band, and white pox disease [11, 37, 38]. Differential exposure to these stressors may result in dissimilar disease resistance by location, with higher occurrences of disease resistance in Florida (27%) relative to similar populations found in Panama (6%) and USVI (8%) [39]. This in turn may influence the prevalence of infection of the nutrient-stress responsive *A. rohweri* [6, 40]. Although disentangling the impact of host and environment will require further sampling and experimental efforts, our comparative analysis of *A. rohweri* populations provides insight into how infection of this symbiont may be influenced by environmental conditions.

Though *A. rohweri* is capable of infecting a variety of non-bilaterian metazoan phyla, the present study focused on infection of *Acropora* coral species found in the Caribbean: *Ac. cervicornis*, *Ac. palmata*, and their hybrid, commonly referred to as *Ac. prolifera*. Our objective was to provide an in-depth analysis of *A. rohweri* population structure and acroporid infections in the Caribbean. We utilized genomic characterization of the three host taxa and their dominant symbiont, *Symbiodinium ‘fitti’*, [32, 41, 42] to investigate possible strain-specific interactions between *A. rohweri* and members of the holobiont. Additionally, we studied the diversity of *A. rohweri* infecting acroporids across the Caribbean to understand how quickly this parasite has evolved in this ecosystem [43–45]. Finally, we compared *A. rohweri* genomes to determine the degree of connectivity between populations and the likelihood of reinfection in the three host taxa.

## Methods

### Sample acquisition and sequencing

Coral samples were collected from across the geographic range of Caribbean acroporids by two research groups. A sample from the Florida Keys was collected as was described in Shaver et al., 2017; this sample was originally used to describe the *A. rohweri* reference genome (Acer44) (GCA 003953955). This sample was extracted and sequenced as described in Klinges et al, 2019. In summary, DNA was extracted using EZNA Tissue DNA Kit (Omega Bio-Tek) and the quality of the extraction was evaluated using the dsDNA HS assay using a Qubit 3.0 Fluorometer. Libraries were prepared using Nextera XT (Illumina) and sequenced using the v3 reagent kit on an Illumina Miseq at the OSU’s Center for Genome Research and Biotechnology (CGRB).

All of the samples presented in Kitchen et al., 2019 and 2020 were evaluated for the presence of *A. rohweri*. Coral tissue samples collected by the Baums’ laboratory originated from twelve reefs across the Caribbean, for a total of 23 *Ac. cervicornis*, 30 *Ac. palmata*, and 23 *Ac. prolifera*. Samples included in this study were collected between 2001 and 2017 (Table S1) and were extracted as described in Kitchen et al., 2019 and Reich et al., 2020, and are briefly summarized here. Sample DNA was extracted using the DNeasy kit (Qiagen, Valencia, CA) and assessed using gel electrophoresis and Qubit 2.0 fluorometry (Thermo Fisher, Waltham, MA) prior to library construction and sequencing by the Pennsylvania State University Genomics Core Facility. A single library from these samples (Acerv FL 14120) was prepared using the TruSeq DNA PCR-Free kit (Illumina, San Diego, CA) and the remaining samples were prepared using the TruSeq DNA Nano kit (Illumina). All Baums’ laboratory samples were sequenced using the Illumina HiSeq 2500 Rapid Run (Illumina, San Diego, CA) and were deposited under PRJNA473816.

### Identifying samples infected with A. rohweri

Sequences were filtered using BBMap version 36.20 to ensure sequences had a minimum length of 30 and a minimum average quality of 10. Reads were aligned to the reference genome (Acer44), using bowtie2 to quantify the proportion of reads that matched the reference sequence and identify candidates for assembly. The possible impact of host and location on the proportion of reads identified as Acer44 were evaluated using two-way analysis of variance (ANOVA) and post-hoc analysis using Tukey HDS within R [46]. The impact of reef location was also evaluated as a nested factor using one-way ANOVA and Tukey HDS. All samples with greater than 10,000 reads matching the Acer44 were *de novo* assembled using Spades version 3.13.1 with the single-cell option to account for possible PCR bias in assembly [47]. Contigs containing *A. rohweri* genes were identified using BLAST and binned as potential genomes; any contigs ≤ 200 bp were excluded from analysis. Genomes greater than 1.2 Mbp were evaluated using CheckM [48, 49].

Metagenome-assembled genomes (MAGs) were compared to the reference (Acer44) and to one another to evaluate commonalities and differences between samples. All samples were compared to one another to establish if there were additional species using pairwise average nucleotide identity (ANI) analysis using OrthoANI [48]. All *A. rohweri* genomes were annotated using Prokka (version 1.14.6) [50] and orthologs common to all samples were identified using orthofinder version 2.3.9 [51, 52]. 1528 orthologs were used to construct a pangenome and identify location-specific orthologs were plotted using the R-program upsetR [46]. Genes identified as being location-specific were further evaluated using the blastx search of the NCBI nr database limited to Rickettsiales [53].

### Evaluating population structure and evolution of A. rohweri

Thirteen additional *A. rohweri* genomes plus the reference Acer44 taken from nine reefs across the Caribbean were used to evaluate potential differences in gene evolution. Phylogenomic trees were constructed with other well-characterized parasitic Rickettsiales species serving as outgroups. Rickettsiales species were selected on the criterium as being closely related to *A. rohweri* and having multiple completed strains with low percent contamination (Table S7); this analysis was performed both to characterize *A. rohweri* evolution and to identify possible shared functions. Rickettsiales genomes were annotated using Prokka (version 1.14.6) and orthologs were identified using orthofinder version 2.3.9 [51, 52]. A total of 143 single-copy orthologs were common to all samples. DNA sequences of orthologous genes were used to generate a phylogenomic tree. This tree was used both as the input for codiversification analysis as well as set the parameters for both evolutionary analysis and the root of the simplified and SNPs phylogenies. The DNA of each individual orthologous gene was aligned using MAFFT version v7.453 [54]. Genes were concatenated by sample and a tree was constructed using IQ-TREE [55, 56] with 1000 bootstrap replicates. Within IQ-Tree, J-modelTest determined the most likely model was the general time reversible model with empirical base and codon frequencies, allowing for a proportion of invariable sites, and a discrete Gamma model with default four rate categories (GTR+F+I+G4).

Whole-genome phylogenetic trends were also evaluated by identifying single-nucleotide polymorphism (SNP) relative to the reference genome *A. rohweri* Acer44. SNPs common to two or more samples that could be used to construct a SNPs phylogeny were found using the haploid SNP-caller, snippy [57], which implements bwa mem and freebayes to identify high-quality SNPs and assemble a core genome alignment. IQ-Tree was used to construct a tree from core-SNPs with 1000 bootstrap replicates; the model selected was jukes canter and ascertainment bias correction (JC+ASC). SNPs impacting annotated portions of the genome were annotated using SnpEFF version 4.3t to find the likely effects of variants on gene function.

Population structure and strain evolution were identified using both inter- and intra-sample diversity. SNPs were found by aligning sample reads to the prokka annotated reference genome (Acer44) using bwa mem, with samclip processing allowing a maximum of a 10 clip length and the removal of duplicate alignments using samtools markdup. To avoid biases that effect variant detection, data was subsampled to the lowest number of reads (2×10^4^) before realigning and then subsampled by the lowest median coverage found in a sample (3), similar to the normalizing protocol outlined in Romero Picazo et al., 2019 [24]. Variants were identified using lofreq with a minimum coverage of 10, a strand multiple testing correlation of p> 0.001, and a minimum SNP quality of 70 were used to find the fixation index (Fst) using the R packages seqinr [24, 58]. Fst estimates the degree of genetic variability between populations, where Fst closer to zero implies populations are mixing freely and Fst closer to 1 implies populations are genetically isolated. SNPs in functional regions were also used to find the intra-sample nucleotide diversity (π), which proves an estimate of the differences in genome diversity between samples. Scripts used to for F_ST_ and π analysis were are publicly available on github repositories (https://github.com/deropi/BathyBrooksiSymbionts). Differences between pairwise F_ST_ values were found within locations within host species was found using one-way with Tukey correction.

Trends in the evolution of functional genes in the *A. rohweri* genome as well as individual genes were evaluated using codeml to estimate ratio of nonsynonymous and synonymous substitutions (dN/dS). Lower values of dN/dS indicating stronger positive selection, that is selection against deleterious mutations to maintain function [59]. Values of dN/dS approaching 1 indicate positive selection or the selection of new mutations into the population. Individual orthologous proteins were aligned using MAFFT, and codon alignments were generated using pal2nal. Codon alignments were evaluated by codeml using a pairwise comparison, with all other parameters set to approximate the tree-building protocol described above. The result of this analysis was compared to the default parameters for codeml. Because the default parameters were less likely than our model parameters, only the results of our model parameters are presented. Only genes with a dN > 0, dS between 0.1 and 2, and dN/dS > 10 were used to find the average dN/dS. This is because pairwise comparisons where dS < 0.01 or dN = 0 are similar enough to be considered identical and dS > 2 are indicative that synonymous substitutions are near saturation. Similarly, values of dN/dS > 10 are considered largely artifactual [60]. From the single-copy orthologs identified by orthofinder, 9% of the total pairwise comparisons are suitable for dN/dS analysis. Within-species and location comparisons were evaluated using two-way analysis of variance (ANOVA) and post-hock analysis using Tukey HDS within R [46].

### Evaluating the impact of environment on A. rohweri replication rate

Because *A. rohweri* has been found in increased prevalence in response to nutrients, the *in-situ* replication rate is likely to be indicative of areas of sustained exposure to nutrients. Replication rate can be estimated for draft-quality genomes using the index of replication, iRep. iRep estimates the raw rate of replication of *A. rohweri* MAGs by realigning reads to the newly constructed genomes. Replication is estimated using an algorithum that assumes bi-directional replication from a single origin and accounts for the total change in coverage in genome fragments. A population where all cells are actively replicating would have an iRep of 2, and a population where only a quarter of the cells were replicating would have an iRep of 1.25 [61]. All genomes contructed had adaquit coverage to perform iRep analysis, but lacked the criterium to produce a filtered result (Table S8); the unfiltered result is presented as the genomes constructed are nearly identical (>99% ANI) and no not necessitate the filtering step to account for strain variation and/or integrated phage that can result in highly variable coverage [61]. Differences in the unfiltered rate of replication as a result of host or location sampled were evaluated using two-way ANOVA and the impact of location was evaluated as a nested factor using one-way ANOVA. All were evaluated using Tukey HDS post-hoc analysis within R [46].

### Identifying likelihood of codiversification of *A. rohweri* with coral or algal symbiont

To evaluate the various possible methods of transmission, multiple codiversification analyses were performed to compare *A. rohweri* evolution with the two common eukaryotic members of the holobiont, *Acropora* and the algal symbiont, the Symbiodiniaceae*, Symbiodinium ‘fitti’*. Both gene-based and SNP-phylogenies were constructed. Coral phylogenies were constructed using mitochondrial control region [62] and full mitochondrial genomes used to identify the parentage of the hybrid species *Acropora prolifera*. The mitochondrial genomes for each individual were assembled using two approaches. In the first approach, filtered and trimmed short-read sequences were mapped to the *A. digitifera* mitochondrial genome sequence (NCBI: NC_022830.1, Shinzato et al. (2011)) using Bowtie 2 v2.3.4.1 (Langmead et al., 2012) with the parameters for the --sensitive mode. Reads were extracted using bamtofastq in bedtools v2.26.0 (Quinlan et al., 2010) and then assembled using SPAdes v3.10.1 (Bankevich et al., 2012) with various kmer sizes (-k 21, 33, 55, 77 and 99). In the second approach, we used the *de novo* organelle genome assembler NOVOplasty (Dierckxsens et al., 2016). The *A. digitifera* mitochondrial genome was used as the seed sequence to extract similar sequences from the original, unfiltered reads for each individual. Consensus sequences of the mitochondrial genomes for each individual were created after manual alignment of the sequences from the two approaches using MEGAX (Kumar et al., 2018). The consensus sequences were run through the web server MITOS (Bernt et al., 2013) to predict genes, tRNAs and rRNAs and non-coding regions. The phylogenetic relationship of the mitochondrial genomes was inferred with the Maximum Likelihood method using RAxML v8.2.12 (Stamatakis, 2014). We included eight acroporid mitochondrial genomes from the Indo-Pacific as outgroups (Liu et al., 2015; van Oppen et al., 2002). Because the mitochondrial genome can undergo different models of evolution among sites, we ran the genome alignment through PartitionFinder v2.1.1 (Lanfear et al., 2016) to determine the best partitioning scheme and substitution models using the greedy algorithm with estimated braches set as linked. In the ML analyses, we used the GTRGAMMA substitution model for the nine partitions. The tree topology with the highest-likelihood based on AIC criterion was reconstructed from 200 replicate trees and nodal support was take from 1,000 bootstrap replicates.

*Symbiodinium* gene trees were constructed using genes described in Pochon et al., 2014 and whole genome SNPs were found as described in Reich et al., 2020, excluding all samples identified as having multiple symbiont infections. All genes were identified using a BLAST search of the aforementioned *de novo* Spades assemblies. Gene trees were aligned using MAFFT and all phylogenies were constructed using IQ-Tree with 1000 bootstrap replicates; genes and model parameters are described in Table S8. Bacterial and eukaryotic phylogenies were evaluated for significant codiversification using the Procrustes Approach to Cophylogeny (PACo), which uses ultrametric rooted trees to create cophenetic matrices that are evaluated for codivergence 10^5^ times in R [63]. This global fit method evaluates phylogenies that are not fully resolved to evaluate if there is significant codiversification between a host and symbiont species. To evaluate transmission by the coral host, bacterial phylogenomic and SNPs trees were compared to coral phylogenies (whole genome SNPs trees and mitochondrial SNPs and genes). To evaluate transmission with a *Symbiodinium* host, bacterial phylogenomic and SNPs phylogenies were compared to *Symbiodinium* gene and SNPs trees. Only comparisons with p< 0.05 are presented along with the residual sum of squares (m^2^) to provide a context for how well the data fit the codiversification model. Significant codiversification analysis were evaluated using a jackknife sum of squares to find those members contributing to the significant association, as they will have values are below the 95% residual sum of squares.

### Evaluating vertical transmission via qPCR of susceptible coral genotypes at early life stages

A few days preceding the annual *Acropora cervicornis* spawning event in 2019 and 2020, sexually mature, adult colonies of genotypes 13 and 50 were brought into Mote Marine Laboratory’s Elizabeth Moore International Center for Coral Reef Research & Restoration from their offshore coral nursery in the lower Florida Keys. On land, corals were isolated to keep replicate colonies and genotypes separated for conducting 2-parent controlled crosses. Spawning, fertilization, settlement, and grow-out were conducted following published protocols and under standardized conditions [64–72]. *A. cervicornis* is a simultaneous hermaphrodite and broadcast spawning activity was observed in August during the predicted peak spawning window for this species [73]. Gamete bundles of eggs and sperm were collected from each genotype, and after bundle dissolution, sperm was isolated from the eggs by filtration using a sieve (100 μm mesh). Triplicate subsamples were concentrated via centrifugation and then snap-frozen and stored at −80°C. Triplicate subsamples of 50-100 eggs per genotype were also snap-frozen and stored at −80°C. The remaining egg stock from genotype 13 and sperm stock from genotype 50 were combined and allowed to undergo fertilization for one hour (cross ‘13e × 50s’). Embryos were reared in replicate cultures with filtered seawater at room temperature (27°C). Approximately one week later, triplicate subsamples of 50 larvae were snap-frozen and stored at −80°C. The remaining larvae were settled in 5-gallon glass tanks using unconditioned ceramic settlement substrates and live, crushed up crustose coralline algae (CCA) as the settlement cue. After settlement, sexual recruits were reared under common garden conditions in Mote’s land-based coral nursery. Corals were maintained in flow-through mesocosms (‘raceways’) with running seawater, from which algal endosymbionts were naturally acquired. Fouling algae was mitigated using intertidal herbivorous snails (*Batillaria spp.),* and the daily husbandry regime consisted of raceway siphoning and basting of the substrates to remove snail detritus. Flow rates were maintained between 4 and 6 L/min depending on season and outdoor weather conditions.

Coral hybrid genotypes were produced from a cross of two disease-susceptible *A. cervicornis* parents (genotypes 13 and 50, Muller et al., 2018) collected from the Mote Marine Laboratory *in situ* coral nursery in 2019 and 2020. The hybrid genotype was sampled at gamete (egg and sperm), larval (<1 week of age), recruit (~2 months), and juvenile (~1 year) stages. DNA was extracted from early life stage coral samples using the Omega E.Z.N.A.^®^ DNA/RNA Isolation Kit. Extracted nucleic acids were stored at −80°C until further processing. Quantitative polymerase chain reaction (qPCR) was performed on 7 early life stage samples (in triplicate) using primers designed to target the *Acropora cervicornis* actin gene (as an endogenous control) and an *Aquarickettsia*-specific gene, *tlc1,* using iQ SYBR Green Supermix (Bio-Rad). An 149bp section of the *tlc1* gene of *Aquarickettsia rohweri* was amplified using sequence-specific primers (F: 5’-GGCACCTATTGTAGTTGCGG-3’, R: 5’-CATCAGCTGCTGCCTTACCT-3’), and the actin gene of *Acropora cervicornis* was amplified as in Wright et al., 2018 [74] as an endogenous control. A sample of *Acropora hyacinthus* (collected from Mo’orea, French Polynesia in 2017) was used as a calibrator, as this species expresses actin but lacks *Aquarickettsia rohweri*. A positive control (a sample of adult genotype 50 with known quantity of *A. rohweri*) and a no-template control (molecular grade water) were prepared using the same methods and quantified simultaneously. A 35-cycle qPCR was performed on an Applied Biosystems 7500 Fast Real-Time PCR System, using cycling parameters selected to minimize mispriming: An initial denaturation step of 3 min at 95°C, followed by 35 cycles of 95°C for 15s, 56°C for 30s, and 72°C for 30s. Melt curve analysis was performed to identify any off-target products. Results were confirmed through non-quantitative PCR of the *tlc1* and 16S rRNA genes (515F-806R primer set, Apprill et al., 2015) using AccuStart™ II PCR ToughMix (QuantaBio) and subsequent gel electrophoresis on a 1.5% agarose gel with Invitrogen 100 bp DNA Ladder (ThermoFisher Scientific).

## Results

### Metagenome Assembled Genomes (MAGs) generated from multiple locations and hosts suggest that A. rohweri associates with many Carribean *Acropora*

*Ca. Aquaricketssia rohweri* sequences were found in all *Acropora* specimen collected from across the Caribbean as part of the Kitchen et al. 2019, 2020 and Reich et al. 2020 studies (Table 1). Samples had between 38 and 1,219,071 reads that aligned to the reference genome, *A. rohweri* Acer44 (GCA 003953955) (Table S1). The *A. rohweri* reads were normalized to the total coral host reads as an intrinsic measure of microbial load [75–77]; using this method, reads varied according to coral host and sampling location (Fig. S1), but not source of collection (Tukey adjusted p=0.5; data not shown). *A. rohweri* made up a greater proportion of the total reads in *Ac. cervicornis* samples relative to *Ac. prolifera* (on average, 4x more reads, Tukey adjusted p=0.0002) and *Ac. palmata* (86x more reads, p<0.0001) samples. Additionally, *A. rohweri* made up a greater proportion of the total reads from all samples collected in Belize relative to Curacao (46x more, p= 0.007). Although limited to *Ac. cervicornis* samples, a greater proportion of *A. rohweri* reads were found in Belize samples relative to those collected from Florida (1.6x more, p<0.00001) and the USVI (1.8x more, p=0.017).

**Table 1:** Collection data and genome quality information for *A. rohweri* constructed from PRJNA473816, including the reference genome from Klinges et al., 2019. Coral host taxa include *Ac. cervicornis* (Acer) and *Ac. prolifera* (Aprol). *Symbiodinium ‘fitti’* infection status, whether by a single strain or multiple strains, were evaluated in Reich et al., 2020. Completeness, contamination, length, percent GC, and N50 were found using checkm, genes are the total number of prokka annotations, and the ANI to the reference sequence *A. rohweri* Acer44 was found using orthoANI.

| Sample ID | Location | Reef | Coll date | Symb infect | Comp (%) | Contam (%) | Length (Mbp) | GC. (%) | N50 | Genes | ANI |
| --- | --- | --- | --- | --- | --- | --- | --- | --- | --- | --- | --- |
| Acer44 | Florida | Key Largo | 1-Jun-13 | unknown | 98.9 | 1.20 | 1.28 | 0.28 | 10,860 | 1,321 | NA |
| Acer FL 4923 | Florida | Key Largo | 22-Nov-11 | solo | 97.6 | 0.10 | 1.21 | 0.28 | 4,669 | 1,243 | 99.9 |
| Acer FL 14120 | Florida | Key Largo | 1-Mar-16 | solo | 100.0 | 0.00 | 1.25 | 0.28 | 12,616 | 1,255 | 99.93 |
| Aprol FI 1303 | Florida | DryTortugas | 29-Jul-03 | multi | 97.8 | 1.32 | 1.23 | 0.28 | 3,890 | 1,343 | 99.91 |
| Acer VI 13738 | USVI | Sapphire | 30-Oct-15 | multi | 98.9 | 0.00 | 1.27 | 0.28 | 24,610 | 1,303 | 99.74 |
| Acer VI 13714 | USVI | Hans Lollik 2 | 29-Oct-15 | solo | 96.1 | 1.53 | 1.26 | 0.28 | 3,725 | 1,299 | 99.66 |
| Acer VI 13708 | USVI | Botany 2 | 28-Oct-15 | multi | 98.9 | 0.00 | 1.29 | 0.28 | 45,966 | 1,302 | 99.75 |
| Acer BE 13797 | Belize | Sandbores 2 | 7-Nov-15 | solo | 100.0 | 0.00 | 1.28 | 0.28 | 22,023 | 1,305 | 99.59 |
| Acer BE 13792 | Belize | Sandbores 2 | 7-Nov-15 | solo | 96.7 | 0.28 | 1.26 | 0.28 | 8,464 | 1,295 | 99.61 |
| Aprol BE 13843 | Belize | Glovers Atoll | 8-Nov-15 | solo | 84.3 | 0.75 | 1.21 | 0.28 | 2,686 | 1,288 | 99.42 |
| Acer BE 13827 | Belize | Glovers Atoll | 8-Nov-15 | solo | 100.0 | 0.00 | 1.29 | 0.28 | 17,891 | 1,302 | 99.61 |
| Aprol BE 13778 | Belize | S.Carrie Bow Cay | 5-Nov-15 | solo | 100.0 | 0.00 | 1.29 | 0.28 | 27,657 | 1,208 | 99.65 |
| Acer BE 13786 | Belize | S.Carrie Bow Cay | 6-Nov-15 | multi | 91.2 | 1.95 | 1.25 | 0.28 | 3,035 | 1,215 | 99.58 |
| Acer BE 8867 | Belize | Curfew | 13-Sep-12 | solo | 100.0 | 0.00 | 1.28 | 0.28 | 27,658 | 1,184 | 99.6 |

Only samples of *Ac. cervicornis* and *Ac. prolifera* from Florida, USVI, and Belize contained sufficient reads to construct *A. rohweri* Metagenome Assembled Genomes (MAGs) (deposited under PRJNA666461). These MAGs were used to compare *A. rohweri* from different locations (Belize: 7, Florida: 4, USVI: 3) and different host-taxa (*Ac. cervicornis*: 11, *Ac. prolifera*: 3). Six of the MAGs constructed were less complete than the original *A. rohweri* genome (<98.9% complete), but four from Belize were 100% complete with no contamination and a larger N_50_ value than the reference assembly (Table 1). All *A. rohweri* MAGs were at least 1.21 Mbp and had a >99% Average Nucleotide Identity (ANI) in pairwise comparisons to the reference genome and one another (Table 1; Table S2).

From the assembled bacterial genomes, between 1184 and 1343 genes were annotated per sample and 98.4% of sequences were identified as belonging to 1528 orthogroups. No orthologs were found to be exclusive to either coral host. Greater than 30% of these sequences were identified as single-copy orthologs shared by all samples. Location-specific orthologs were all single-copy; Florida had 8 unique orthologs, Belize 40, and USVI 21 (Fig. 1). The majority of location-specific genes were annotated by prokka as hypothetical proteins; however, searching for the function of these genes against the NCBI nr database resulted in additional annotation. Functional genes specific to locales include: the protein transfer gene *secA* in Florida, as well as multiple transport genes in Belize and USVI, as well as two genes involved in the type II toxin-antitoxin system in Belize (Table S3).

**Fig. 1:**
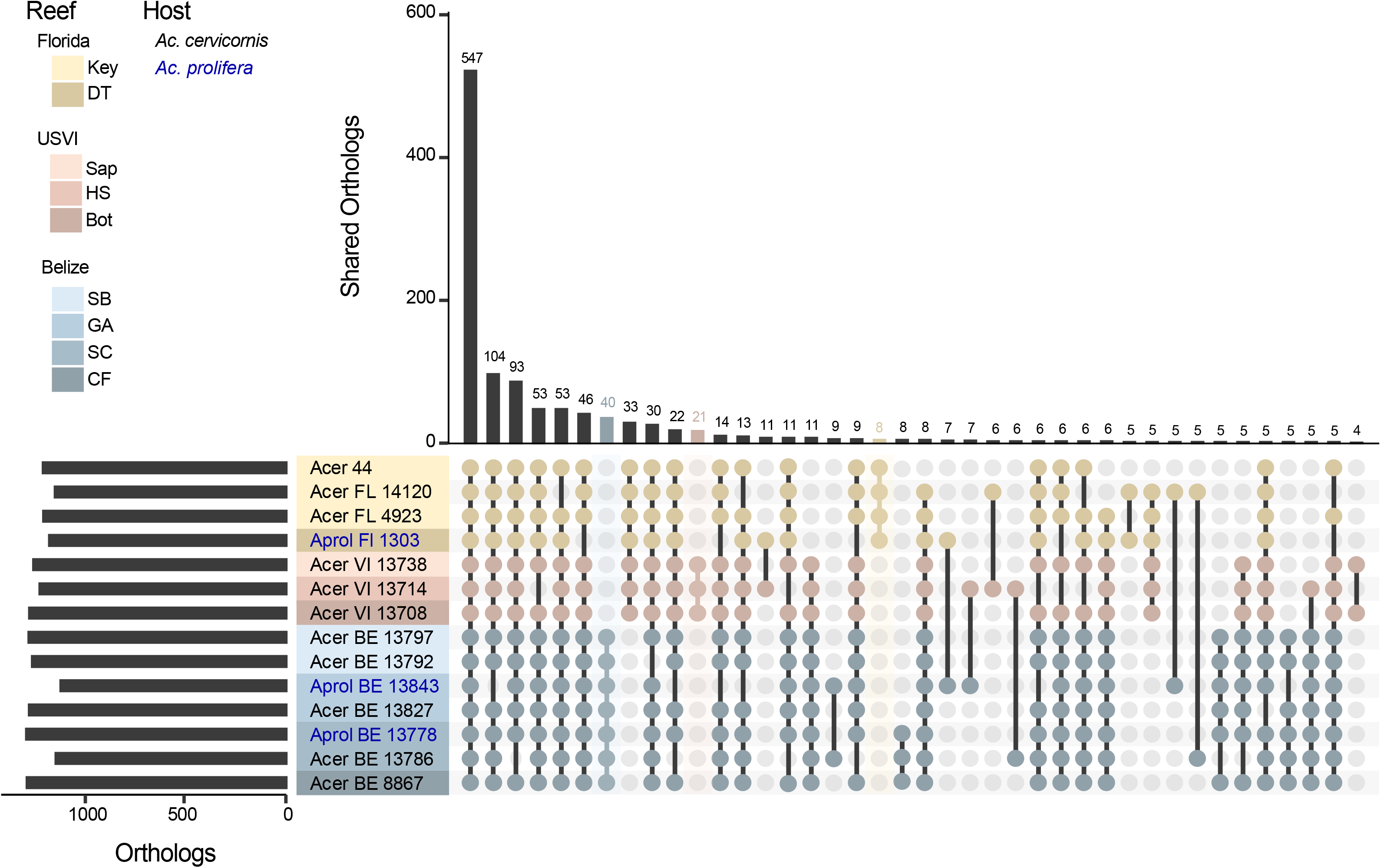
The number of orthologous genes in *A. rohweri* identified using orthofinder for each sample. Location specific genes are highlighted in the bar plot by location color: Florida (yellow, 8 orthologs), USVI (pink, 21 orthologs), and Belize (blue, 40 orthologs). Host identity is noted in blue text for *Ac. prolifera* (Aprol) and black text for *Ac. cervicornis* (Acer), although no orthologs are exclusive to either host taxon.

### The coral parasite, A. rohwerii, differentiates by location, not by host

Phylogenomic analysis of the orthologous genes of the *de novo* assembled *A. rohweri* MAGs showed variation among samples collected from different locations, irrespective of host. Comparison to well-studied host-associated Rickettsiales species (*Ehrlichia chaffeensis*, *Rickettsia prowazekii*, *Rickettsia rickettsii*, and *Wolbachia pipientis*) resulted in 71 orthologous genes. The phylogenetic tree constructed from DNA of these orthologs resulted in all newly constructed *A. rohweri* genomes tightly clustered near the reference genome Acer44 (Fig. S2). Limiting phylogenomic analysis to *A. rohweri* samples results in clear differentiation between samples collected across the Caribbean and north-west Atlantic; Belize *A. rohweri* are distinct from those isolated in the Virgin Islands and Florida, regardless of host identity (Fig. 2A). Further, the *A. rohweri* isolated from *Ac. prolifera* does not form a separate clade from those isolated from *Ac. cervicornis* even when collected from the same location. This is especially evident in Belize samples; *A. rohweri* genomes constructed from *Ac. prolifera* and *Ac. cervicornis* taken from the same reef are more closely related than *A. rohweri* collected from *Ac. prolifera* from a neighboring reef. Clustering by location was similar in the SNP analysis (Fig. 2B), despite there being a surprisingly small number of SNPs per sample (*n*=11 to 2345), at a minimum read depth of 10, minimum fraction of 0.9, and a minimum mapping quality of 100 (Table S4). Relative to the reference genome Acer44, samples had low levels of genetic polymorphism (0.63 ± 0.69 SNPs/kilobase) (Table S4). This resulted in few SNPs that are shared by multiple samples (*n*=15), i.e. those that are informative in phylogenetic analysis.

**Fig. 2:**
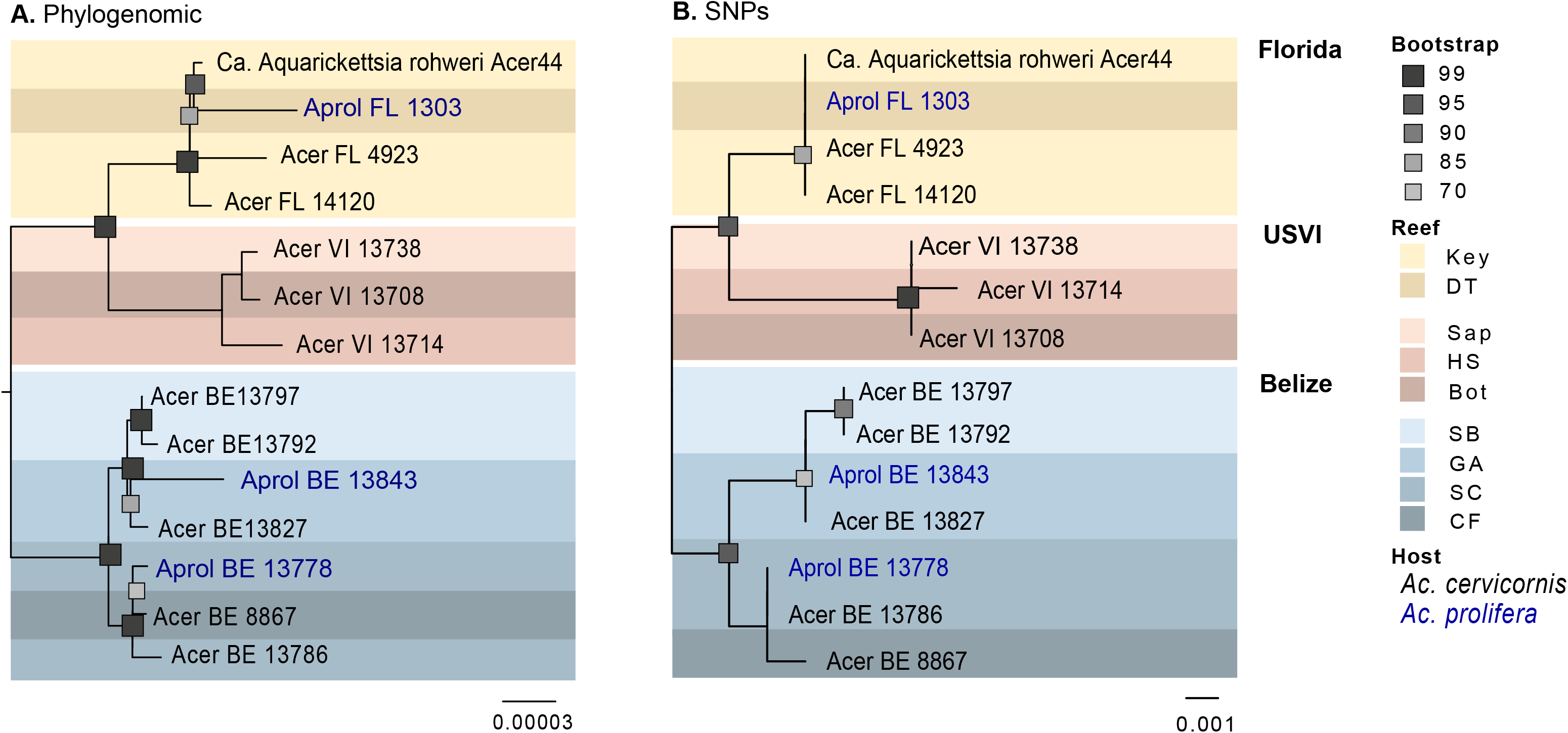
*A. rohweri* phylogenies for all MAG generated using (A) orthofinder to identify orthologs to construct a phylogenomic tree, rooting based on comparison to other Rickettsiales (Fig. S1) (B) Phylogenetic tree of core-SNPs (15) generated by snippy and rooted based on the outcome of orthofinder tree. Bootstrap values greater than 70 are shown.

Analysis of inter- and intrasample SNPs within annotated portions of the genome indicates that although samples have similar levels of genetic diversity, the bacteria found within individual coral colonies are relatively genetically isolated from one another. This is even true of samples collected from the same reef (Fig. 3). Intrasample nucleotide diversity (π) did not differ among sampling locations (reefs) and was on average 1.85 ± 0.8 × 10^5^. However, pairwise comparison of intrasample *Ac. rowheri* SNPs between coral samples mostly resulted in an fixation index (F_ST_) of 0.86 or greater, suggesting *A. rowheri* infections are genetically isolated from one another by coral sample. This level of genetic isolation was found in all pairwise comparisions in USVI and Belize, and was even observed in pairwise comparisions of samples taken from the same reef. Samples from Florida had significantly lower fixation indices relative to USVI and Belize (F_ST_ =0.65-0.83; Tukey adjusted p<0.0001) Although these values still imply *A. rowheri* populations in Florida coral colonies are somewhat genetically isolated from each other, these populations may be less fixed than those found in USVI or Belize. The aforementioned account for *A. rohweri* found in both *Acropora* taxa; host identity did not affect the level of genetic isolation (Tukey adjusted p=0.9).

**Fig. 3:**
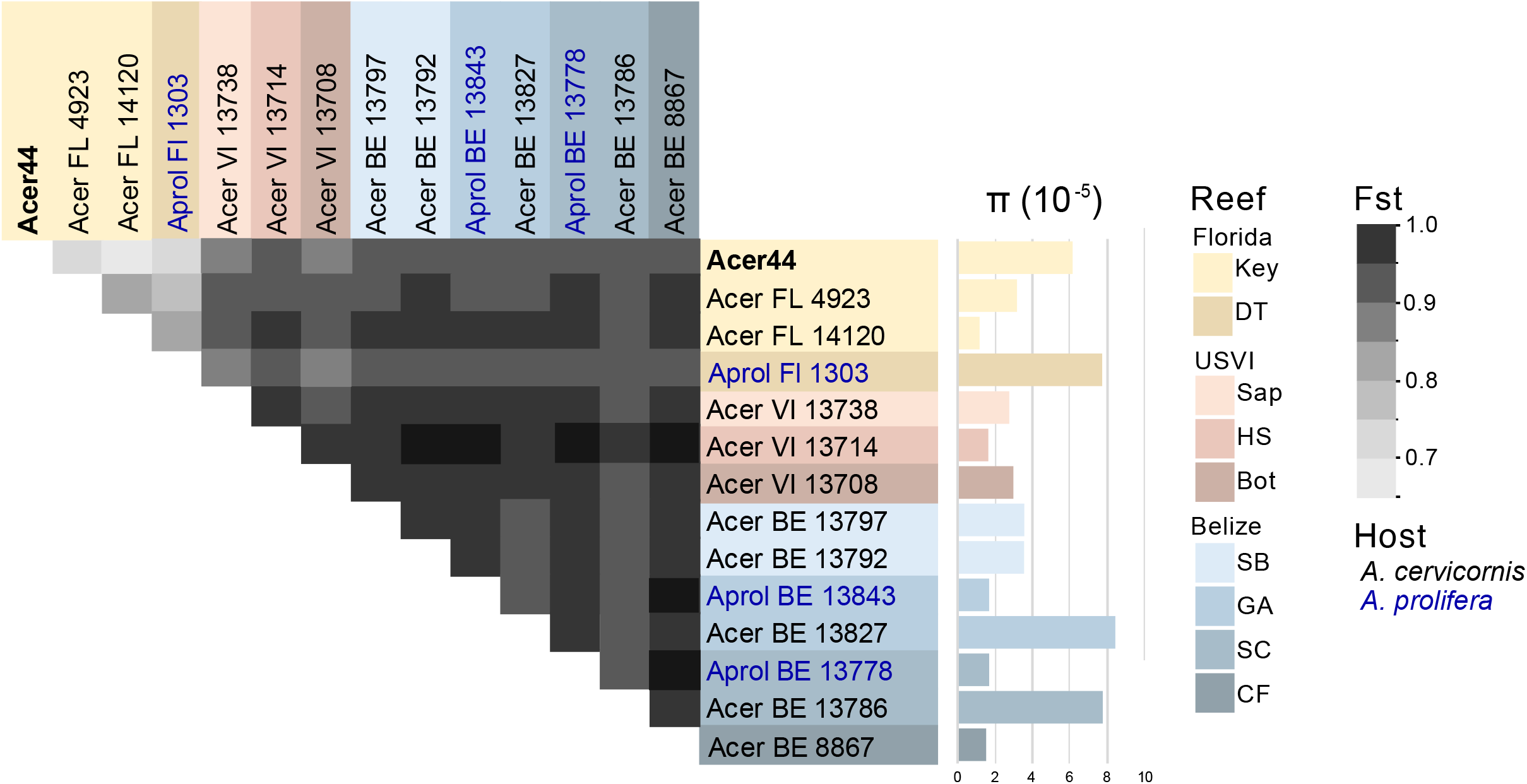
Pairwise comparisons of the fixation indices (F_ST_) and intrasample nucleotide diversity (π) for each sample from each location, generated using methods outlined in Romero Picazo et al., 2019 [24]. Samples originating from Belize shown in blue, Florida in yellow, and the USVI in orange. All comparisons between USVI and Belize samples resulted in an F_ST_ of > 0.86 whereas samples with the Florida populations were between 0.64-0.83.

All samples had high-quality, intersample SNPs identified using snippy [57] with annotated functional genes relative to the reference genome *A. rohweri* Acer44. Of those SNPs that impacted functional genes, a majority of the SNPs (on average 62%) were identified as missense mutations (Table S4). Three samples from Belize (Aprol BE 13843, Acer BE 13797, and Acer BE 13792) were found to have a nonsense stop gain mutation in the IS66 family transposase ISDpr4. Four additional transposases were identified as having missense mutations, including IS6 family transposase ISCca2, which was found to have a missense mutation in all Belize and USVI samples. These SNPs were annotated by SnpEFF as having a moderate impact on function (Table S5).

### A. rohweri undergoing positive selection

Samples were compared to well-studied Rickettsiales relatives of *A. rohweri* to evaluate whether populations of *A. rohweri* are undergoing neutral or positive selection relative to other closely related parasites. *A. rohweri* had the highest mean dN/dS, with a much broader distribution of dN/dS values relative to all other Rickettsiales species (36-89% greater dN/dS, p<0.0001) (Fig. 4A; Table). This indicates *A. rohweri* is undergoing greater positive selection than closely related Rickettsiales. Comparison of dN/dS of *A. rohweri* from the two *Acropora* host taxa did not result in significantly different dN/dS (p=0.06; data not shown), but location did affect dN/dS. There was a higher median dN/dS for *A. rohweri* from Florida relative to those from USVI (65% higher, p=0.048) (Fig. 4B). However, all populations had some genes undergoing positive selection (Table S5). Most of these genes were identified by prokka as hypothetical. Of those that were not hypothetical, GTPase Era, DUF2312 domain-containing protein, and the Bifunctional protein FolD were consistently undergoing some level of positive selection (0.5-0.9 dN/dS) in all samples taken from Belize. The strongest positive selection (≥ 1.0 dN/dS) were 50S ribosomal protein L13 and the type IV secretion system protein VirD4, although selection was only observed in a subset of comparisons between *A. rohweri* samples (Table S7).

**Fig. 4:**
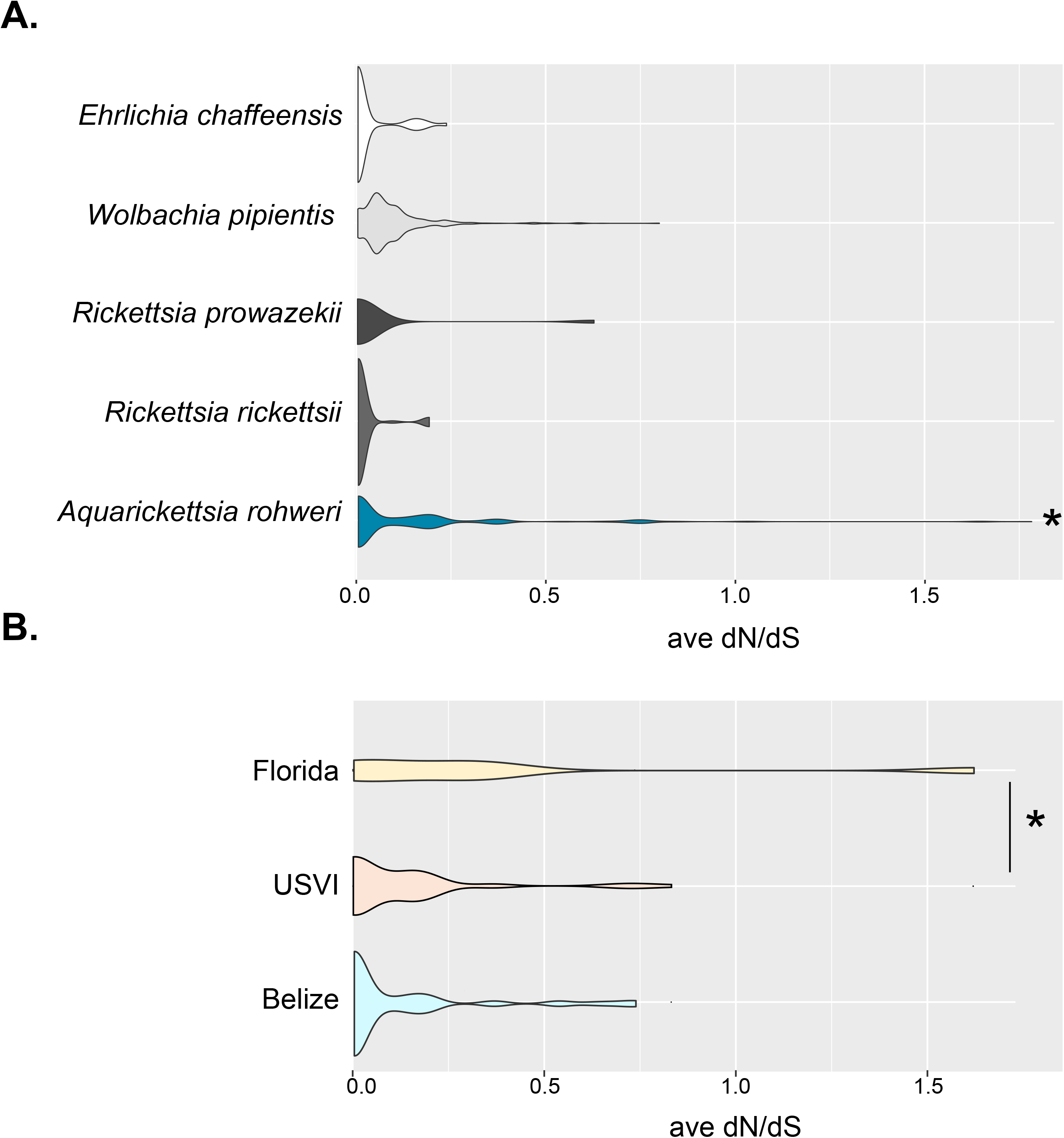
Plots of average dN/dS values for whole genome comparisons of prokka annotated genes. (A) Average dN/dS of closely related well studied Rickettsiales species were all lower relative to *A. rohweri* (p<0.0001). (B) Average dN/dS of *A. rohweri* is significantly greater in Florida than USVI (p =; 0.048).

### A. rohweri from Florida samples had higher replication rates

Samples collected from Florida had consistantly higher estimated unfiltered rates of replication (iRep), with the identity of the coral taxon having no impact (Tukey adjusted, p=0.3) (Fig. 5). The unfiltered iRep of samples from Flordia was 19% higher than samples from Belize and 30% higher than samples from USVI. A single sample from Belize had as high an iRep as Florida samples (Acer BE 13786); this sample was taken alongside five other Belize samples and also from the same reef as an additional sample (Aprol BE 13778 from South Carrie Bow Cay). Florida samples had a significantly higher estimated rate of replication than those taken from the USVI (p=0.04); no difference in rate was seen when comparing iRep among reefs (p=0.44).

**Fig. 5:**
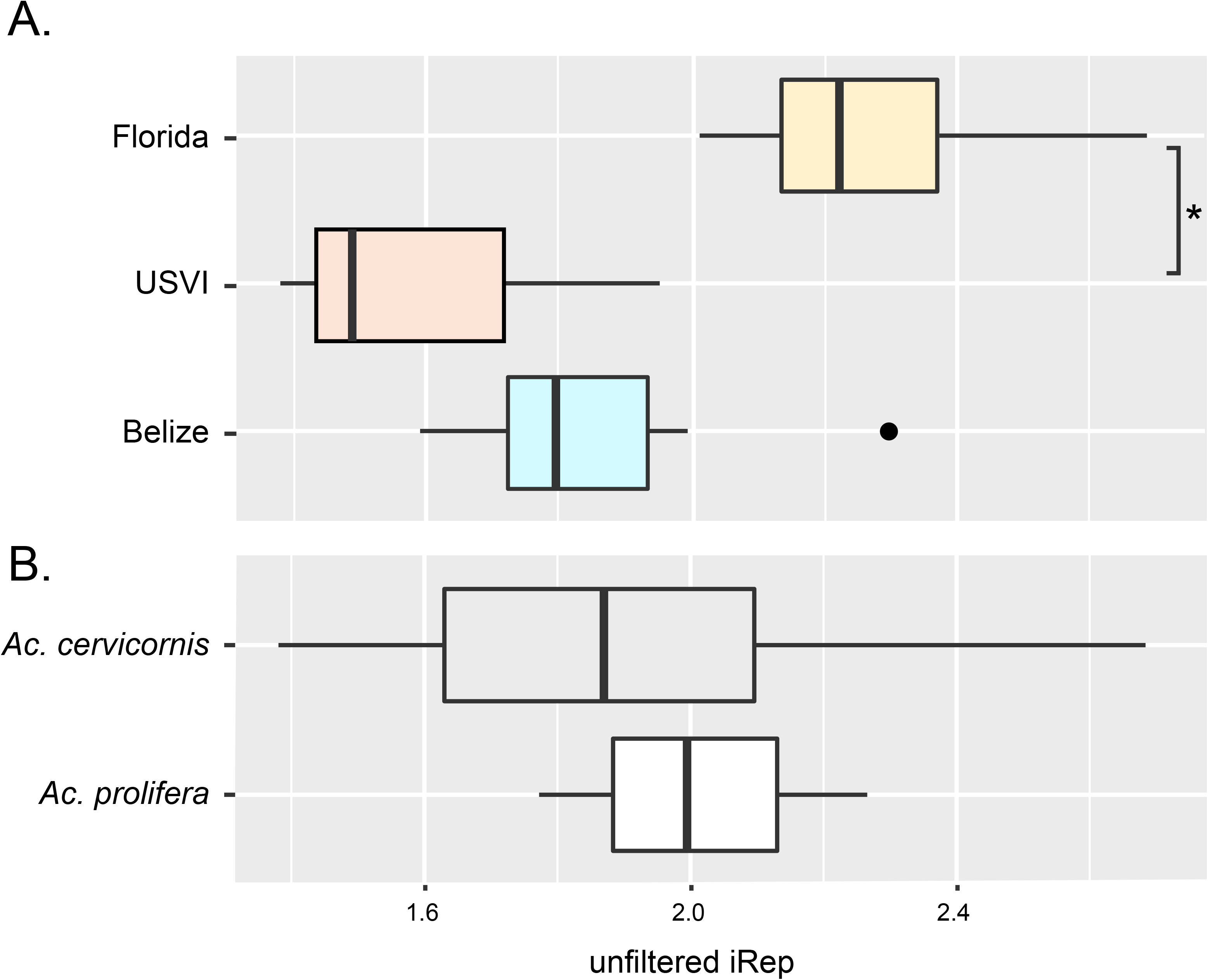
The average unfiltered estimates for the rate of replication for (A) sampling locations and (B) coral host.

### Codiversification analysis and qPCR of coral offspring suggests horizontal transmission

All phylogenetic methods used to analyze the coral diversity of samples infected with *A. rohweri* resulted in differentation between *Ac. cervicornis* and *Ac. prolifera* (Fig. S3, S4). Codiversification analysis among *A. rohweri* and coral phylogenies largely were not signficant. Although coral phylogenies constructed using the mitochondrial control region identified by Vollmer et al., 2002, were signficantly codiversifying with bacterial phylogenomic and SNPs trees (p=0.02), the m^2^ values indicate the data does not fit the codiversification model and are therefore highly unlikely (>800). Coral phylogenies contructed using the whole coral mitochondrial genome also did not codiversify with either *A. rohweri* phylogenomic or SNP phylogenies (p<0.01 and m^2^>330). This was recapitulated in the jackknife sum of squares of both Procrustes Approach to Cophylogeny (PACo) analysis, which did not identify any samples as significantly contributing to codiversification (Table S8).

Analysis of the *S. ‘fitti’* gene trees and *A. rohweri* SNP phylyogenies were also incongruous. The phylogeny constructed using the genes recommended in Pochon et al., 2014 resulted in placement of all samples as *Symbiodinium ‘fitti’* (ITS2 type A3, formerly “Clade A”; Fig. S5). The gene tree also showed a clear differentiation between the samples collected from Florida relative to the USVI and Belize samples. However, USVI and Belize samples formed a single clade, which is dissimilar from the clustering that occurred in either *A. rohweri* phylogeny (Fig. S5). As was previously described in Reich et al., 2020, the *S. ‘fitti’* SNPs phylogeny clustered by host species and not location (Fig. S6). Similar to the coral analysis, codiversification analysis of *S. ‘fitti’* with *A. rohweri* phylogenies were mostly insignificant. Only the *S. ‘fitti’* gene tree had significant codivergence with *A. rohweri* phylogenomic and SNPs phylogenies (p=0.02), however the m^2^ values were also prohibitively high for this analysis (m^2^>380), and jackknife analysis identified 0-2 samples contributing to this trend (Table S7).

Quantitative PCR was utilized to detect *A. rohweri* in early life stages of *Acropora cervicornis*. qPCR was performed on samples of gametes (egg and sperm), larvae (<1 week of age), recruits (~2 months), and juveniles (~1 year) of a hybrid genotype produced by crossing two disease-susceptible *Acropora cervicornis* parents (genotypes 13 and 50, Muller et al., 2018). A calibrator sample (*Acropora hyacinthus*, known to lack *A. rohweri*) and a positive control (adult genotype 50 *A. cervicornis*, with an average relative abundance of *A. rohweri* greater than 70%) were also quantified. While adult *A. cervicornis* collected from the Mote *in-situ* coral nursery had high fold change of *tlc1* relative to the calibrator sample, amplification of *tlc1* across all early life stage samples was not significantly different from the calibrator sample and led to the failure of the thresholding algorithm, indicating an absence or undetectable level of *A. rohweri* infection in these early stages of *A. cervicornis* ontogeny.

## Discussion

*Ca.* Aquaricketssia rohweri infection was found in every sample of acroporid corals taken from across their Caribbean and north-west Atlantic geographic range, with *Ac. cervicornis* samples consistently yielding more *A. rowheri* reads relative to *Ac. prolifera* and *Ac. palmata* (Fig S1A). This coincides with relatively higher disease susceptiblity of *Ac. cerivornis* [6, 8, 9]. Assuming the proportion of *A. rohweri* to host reads are indeed reflective of infection status [75–77], relative resistance of *Ac. palmata* may be attributed to environmental factors such as depth, innate host immunity, or defenses mounted by the host microbiome [78–80]. Determining which factors may be leading to resistance in *Ac. palmata* is a valuable area of further research.

The coral *Ac. cervicornis* yielded higher read numbers of *A. rohweri*, but both *Ac. cervicornis* and *Ac. prolifera* hosted sufficient reads to construct *A. rohweri* genomes of similar length and quality as the reference genome Acer44. Phylogenomic analysis using orthologous genes and SNPs indicate the bacteria infecting Caribbean acroporids are specific to the collection location and not the host. This is in contrast to the only other acroporid symbiont with population genetic information, the dinoflagellate *S. fitti*. Strains of *S. fitti* preferentially associate with each host, regardless of reef location. These contrasting patterns of population structure indicate that the forces shaping the two coral symbionts are different despite their shared host. Differing population structures have also been reported in terrestrial symbionts, including those populating aphids and whiteflies [27, 81, 82].

*A. rohweri* populations from the three sampling locations form separate clades in phylogenomic and SNP phylogenies, with Florida and USVI samples likely sharing a closer ancestral lineage than Florida and Belize. However, USVI is not basal to Florida populations, suggesting that *A. rohweri* from Florida did not originate from the US Virgin Islands. A barrier to gene flow has been identified between the eastern and western Caribbean for the coral host [83, 84], suggesting there is a barrier to planktonic genflow between these environments. Similar genetic differentiation by location has been observed for the *Ac. cervicornis* sequences of the same samples used in this analysis [41]. Even though seasonally variable surface currents connect all sampling locations [84, 85], and all samples are genetically similar (relative to the reference genome, all samples were >99% ANI), the differences found reveal clear differentiation between Florida, the US Virgin Islands, and Belize *A. rohweri* populations.

*A. rohweri* collected across the Caribbean have low levels of genetic polymorphism with <2500 SNPs relative to reference genome of 1.28 Mbp. Thus, *A. rohweri* may be considered monomorphic (*sensu*, 54-56). Although the rate of genetic polymorphisms in *A. rohweri* is similar to that of bioluminescent mutualists *Ca*. Photodemus katoptron and *Ca*. Photodemus blepaharus, it is also comparable to that seen in pathogens *Yersinia pestis* and *Bacillus anthracis*, [86–88]. Lower levels of genetic polymorphism are correlated with virulence in some bacteria [88]. The role of *A. rohweri* in coral disease is an active area of research, and thus it is difficult to interpret the low levels of genetic polymorphisms in *A. rohweri*, but it is notably low for a symbiont spanning such a large geographic range.

Of the subset of SNPs that impact functional regions, most (62%) resulted in a change to the amino acid identity and therefore likely affect protein function. Although the majority of genes impacted by missense mutations were hypothetical proteins, some gene annotations were identified as transposases. Moreover, the single gene found to have acquired a stop codon in all USVI and Belize samples was within the transposase ISDpr4. The loss of transposases and the eventual loss of these gene regions is characteristic of long-term obligate symbionts [16, 89]; therefore, the *A. rohweri* genome may still be actively reducing through the loss of mobile genetic elements.

Although the bacteria were closely related to each other and are phylogenetically clustering by sampling location, there is surprisingly little genetic mixing, even within a single reef. Pairwise comparisons of the fixation indices between all samples had such high values that genetic mixing among samples and thus reinfection is unlikely to occur between or even within a reef. The most extreme case was observed in comparing samples collected from the same reef in Belize, as all pairwise comparisions had an F_ST_ of 0.95 or greater. The lowest level of pairwise genetic differences were observed in Florida, which implies a slightly higher probability of reinfection relative to the other populations sampled, but laboratory studies would be needed to evaluate whether this is due to host or environmental factors. Thus, despite low genetic diversity observed overall, genetic diversity was distributed such that locations and samples were highly differentiated suggesting that *A. rohweri* infection may occur earlier in the coral lifespan and propagate within the host with little to no genetic mixing occuring thereafter.

Our work also shows that *A. rohweri* is undergoing greater positive selection relative to closely related parasitic Rickettsiales species, with genes involved in speciating and virulence being most impacted. While the identity of the coral host did not matter, sampling location did impact did impact the degree of positive selection. The average dN/dS of Florida samples is, on average, 2.7 times greater than samples from USVI and 2.8 times greater than those in Belize. Although differences in dN/dS were not observed for all samples at each location, these trends were observed in a subset of the comparisons between all sampling locations. Genes that were associated with ribosomal assembly, L13 and GTPase ERA, which assemble 50S and 30S ribosomal proteins respectively, were undergoing positive selection in a subset of the samples. The consequence of ribosomal-associated genes undergoing positive selection is unknown, but it may be indicative of speciation occurring between the different sampling locations. Another gene undergoing positive selection in a subset of samples across locations was the Type IV secretion system-coupling protein VirD4. VirD is essential to T4SS and is involved in substrate recruitment, which plays a role in oncogenic DNA transfer and virulence in Agrobacterium [90–93]. Thus, positive selection in VirD may be affecting how *A. rohweri* in Florida populations interact with their host species.

Though microscopy found *A. rohweri* living in close proximity to coral and S. *‘fitti’*, neither are likely transmitting the parasite vertically. Coral larvae seemed the most likely method for transmission across the Caribbean, as larvae can travel long distances as plankton (>500 km) [84]. Similarly, algal symbionts could provide the necessary nutrients to *A. rohweri* and facilitate parasitic infection when *S. ‘fitti’* is acquired by juvenile coral hosts [94–97]. It is also possible that the parasite could be carried alongside either member of the coral holobiont as they reproduce asexually. However, either sexual or asexual reproduction would have resulted in congruence between the parasite and host phylogeny. Yet, codiversification analysis of both SNPs and gene-based phylogenies resulted in neither coral nor *S. ‘fitti’* having clear codivergence with *A. rohweri*. Additionally, qPCR evaluation of early life phases (<1 week to 1 year) from disease-susceptible coral genotypes known to harbor *A. rohweri* as adults did not have detectable *A. rohweri*. The reduced metabolic capabilities of *A. rohweri* [1] and the lack of evidence for a dormancy pathway also suggests that the bacteria is unlikely to survive long periods in the environment as free-living bacteria. It is therefore most likely that the bacteria are transmitted *via* an alternative method that would provide the necessary nutrients, such as through the movement of coral mucocytes coupled with some abrasion or inoculation event or through an as yet unidentified intermediate host.

Overall, the results of this study show that *A. rohweri* infection differs among coral hosts and locations, is evolving at different rates across its host’s range, and is horizontally transmitted. These findings suggest new pathways to the study of *A. rohweri* and its potential contribution to coral diseases in the Caribbean. For example, exploring possible host or microbiome-based deterrents of *A. rohweri* infection of *Ac. palmata* [98] may be valuable to the preservation of Caribbean acroporids. Additionally, Florida may be a unique focal point for the study of how *A. rohweri* infection impacts coral disease progression. Several *Ac. cervicornis* and an *Ac. prolifera* from the Florida Keys host high concentrations of *A. rohweri* that tend to be less isolated, undergo greater selection in speciation and virulence genes, and are propagating faster in Florida than in other sampling locations. Thus, further research into environmental stressors and host responses in this population will be invaluable to our understanding pathogen evolution, its role in coral disease, and the restoration and recovery of this fragile ecosystem.

## Supporting information

all supplemental materials

## Data accessibility

All sequences used are available in the SRA in PRJNA473816, assembled *A. rohweri* genomes are accessible under PRJNA66646, and coral mitochondrial genomes are MW246489-MW246565.

## Acknowledgements

Thanks to Dr. Tory Hendry for feedback on evolutionary methods employed and Dr. Hanna Koch for permits facilitating the movement of corals between Mote’s field and land coral nurseries; Florida Keys National Marine Sanctuary under permit # FKNMS-2015-163-A3.

