## Supplementary figures and images for "The coral symbiont *Candidatus* Aquarickettsia is variably abundant in threatened Caribbean acroporids and transmitted horizontally"

### Fig. S1_Percent A.rohweri reads data.pdf

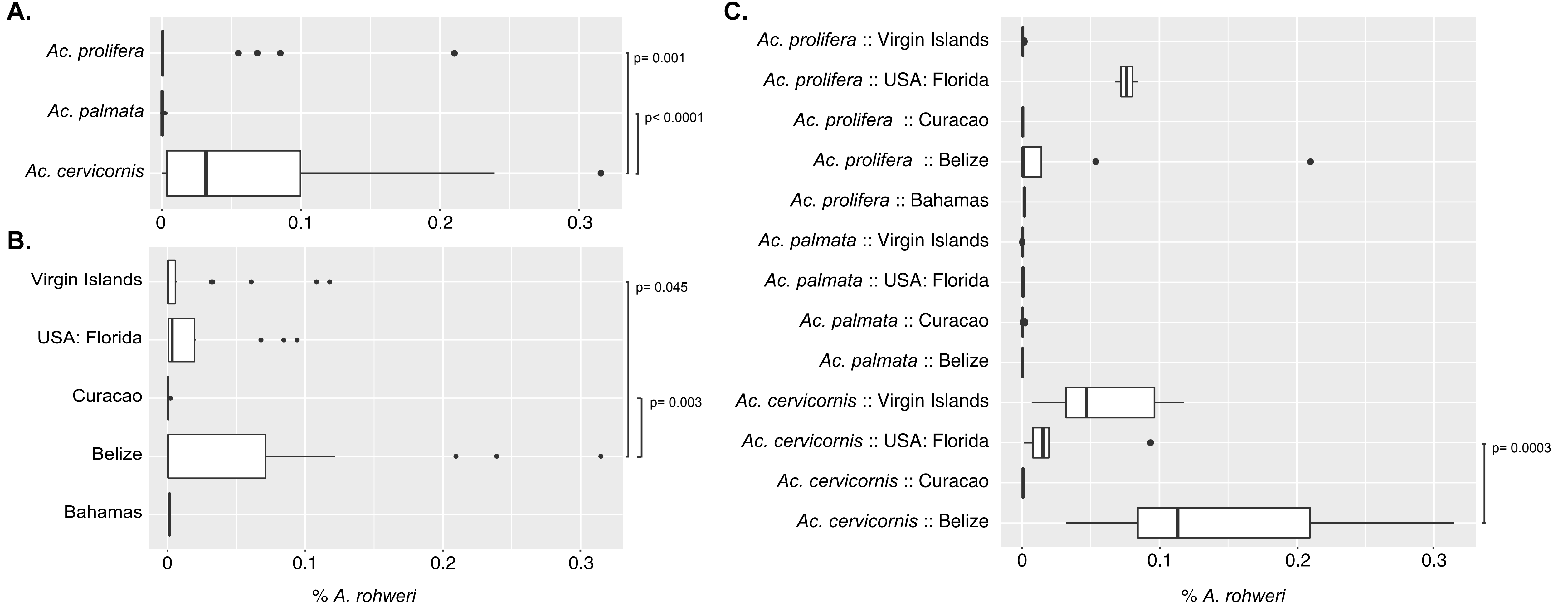

### Fig. S1_Percent A.rohweri reads data_redo.pdf

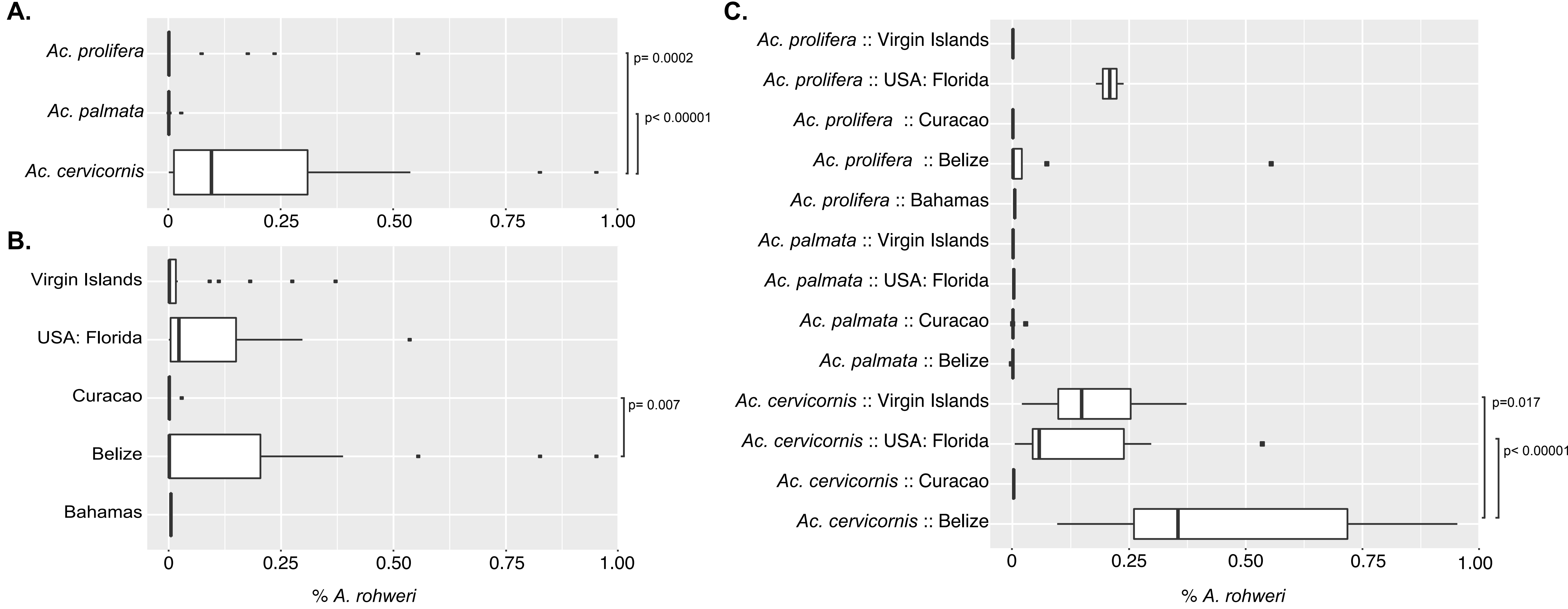

### Fig. S2_Bacterial phylogenomic tree.pdf

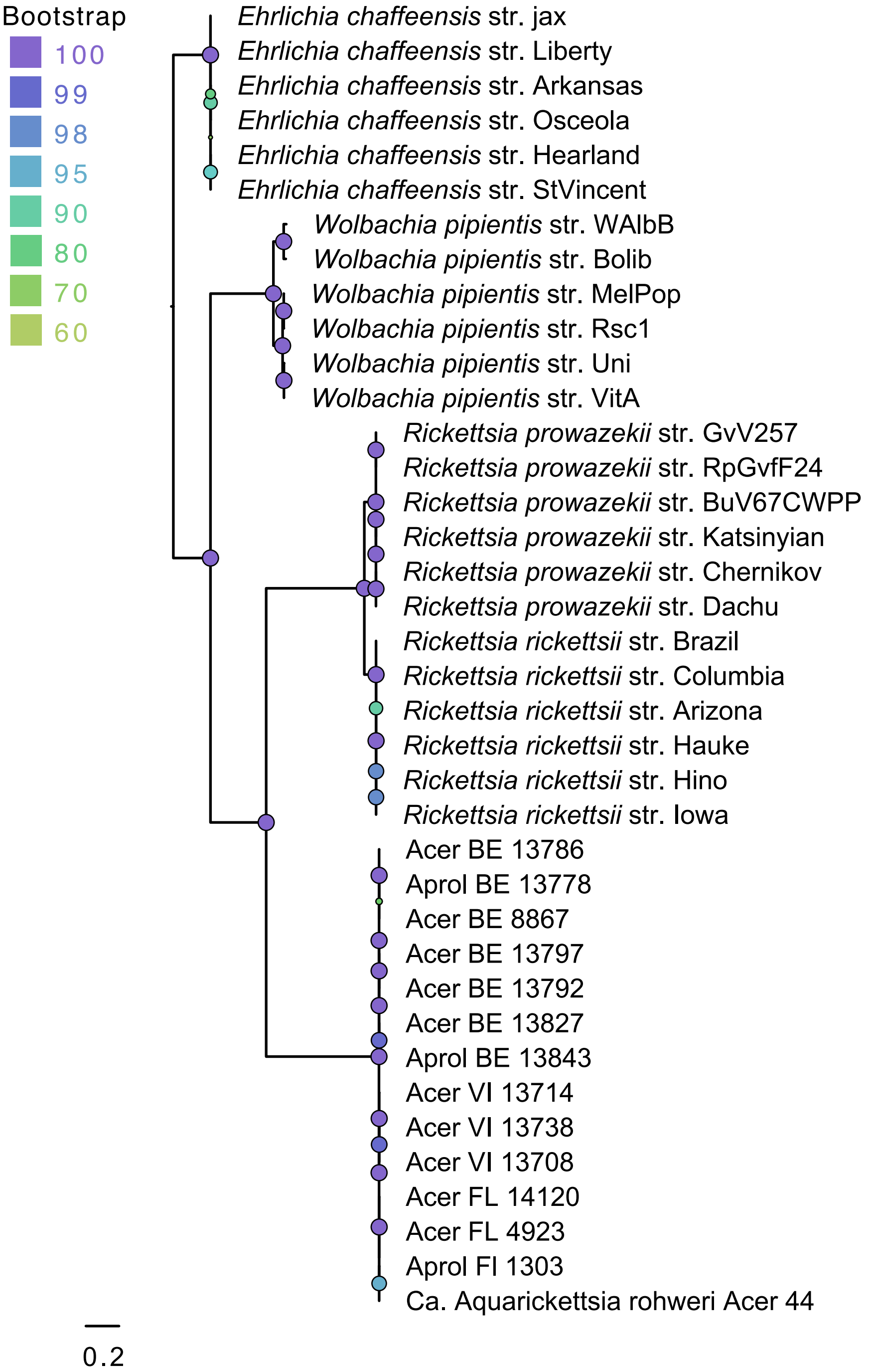

### Fig. S3_Coral mitochondrial control region.pdf

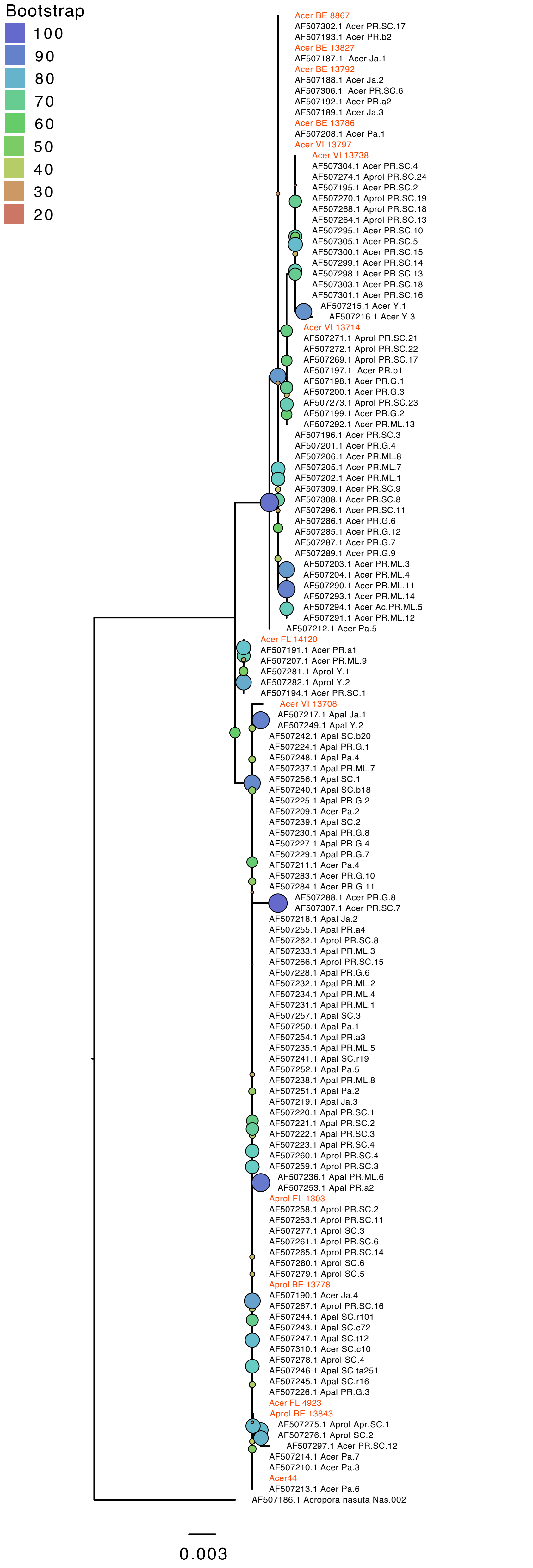

### Fig. S4_Coral_mitoSNPs_ultrametric.pdf

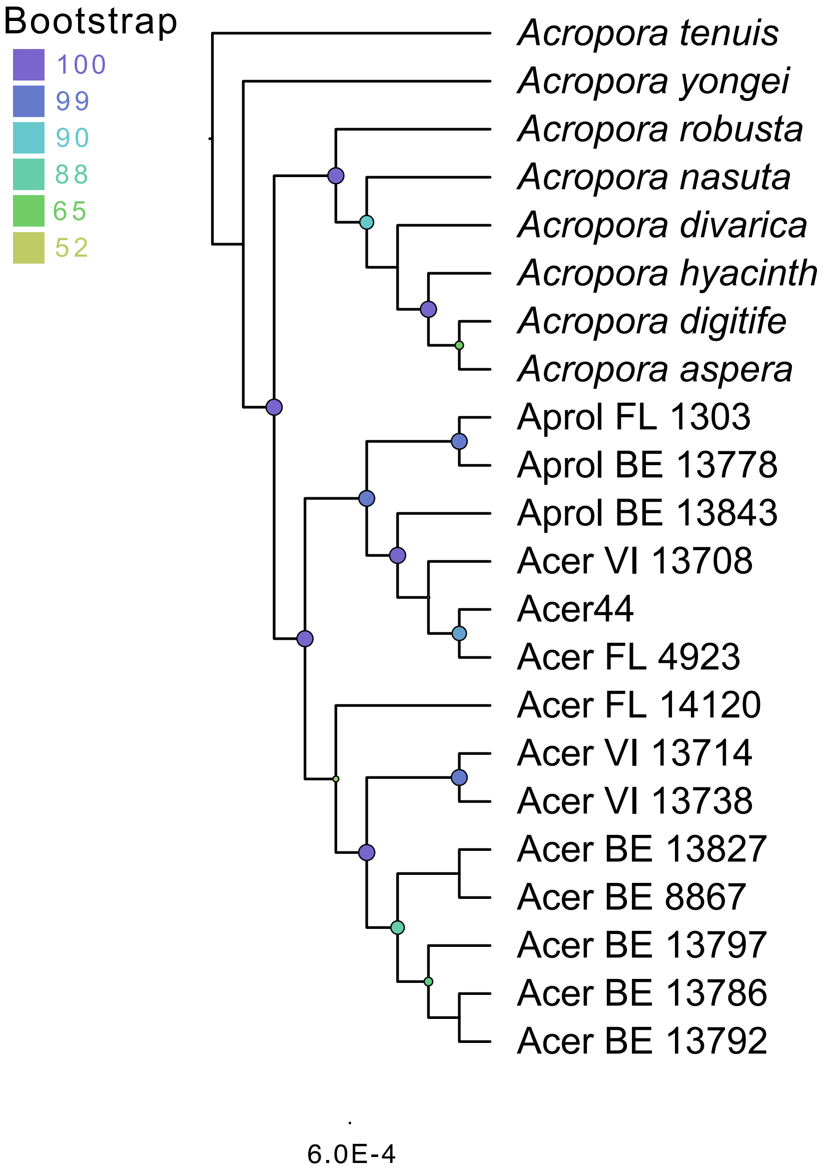

### Fig. S5_Symbiodinium_genes_ultrametric.pdf

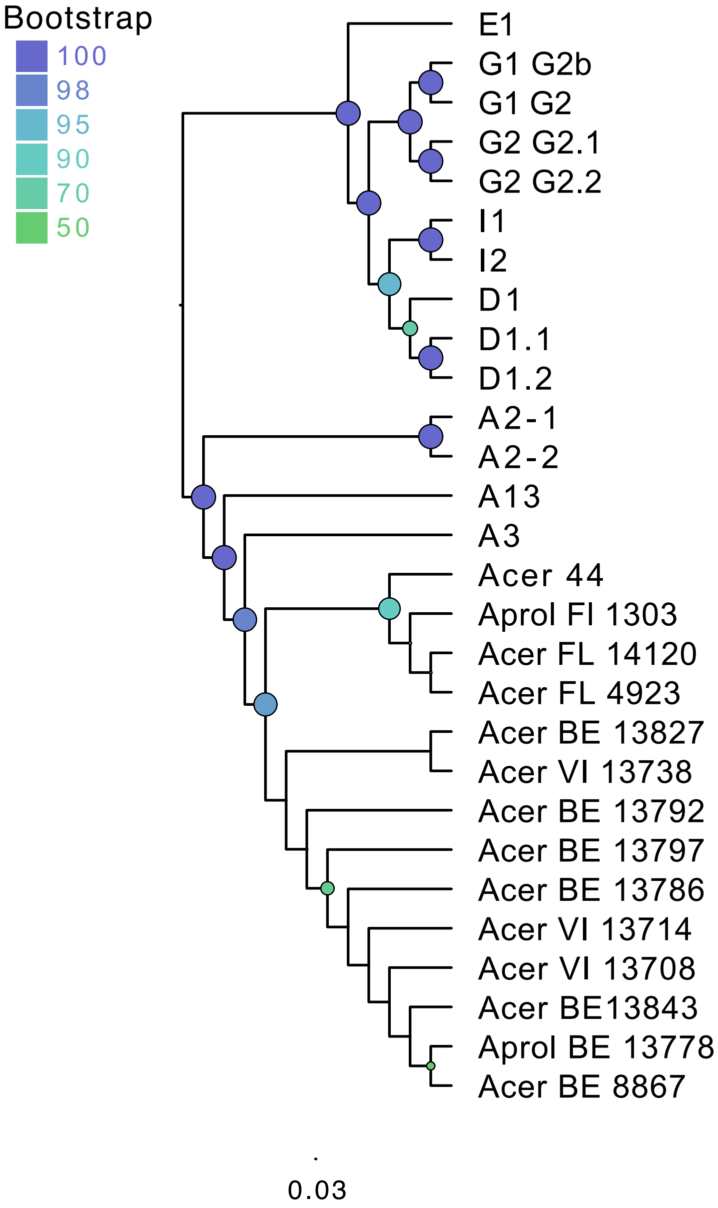

### Fig. S6_Symbiodinium snps._tree.pdf

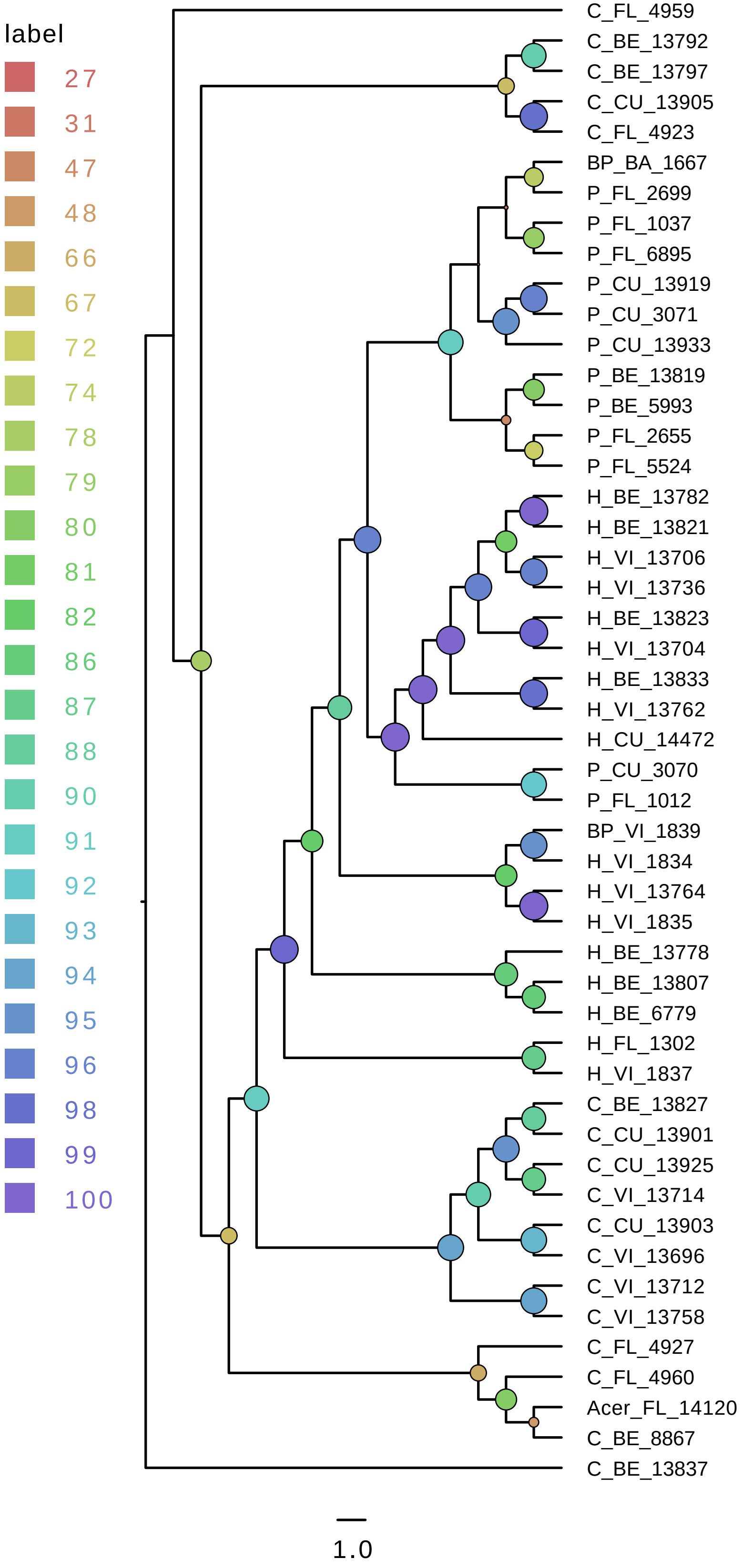
